# Rice Annotation Project Database (RAP-DB): literature-curated gene annotation and integrated omics resources for rice functional genomics and molecular breeding

**DOI:** 10.64898/2026.01.16.699882

**Authors:** Yoshihiro Kawahara, Tomoko Hirozane-Kishikawa, Ryo Hirata, Xiaohui Wang, Yuki Tamagaki, Masahiko Kumagai, Norio Tabei, Hiroaki Sakai, Takeshi Itoh

**Affiliations:** Research Center for Advanced Analysis, National Agriculture and Food Research Organization (NARO), Ibaraki 305-8602, Japan; IMSBIO Co., Ltd., Tokyo 170-0013, Japan; Mizuho Research & Technologies, Ltd., Tokyo 101-8443, Japan; DYNACOM Co., Ltd., Chiba 261-7125, Japan; Master Program in Global Agriculture Technology and Genomic Science, International College, National Taiwan University, Taipei 10617, Taiwan; Center for Computational and Systems Biology, National Taiwan University, Taipei 10617, Taiwan

**Keywords:** Rice, Gene annotation, Literature-based manual curation, Artificial intelligence, Transcriptome, Agronomically important genes, Genome diversity, Allelic variation

## Abstract

High-throughput sequencing technologies have enabled the generation of high-quality reference genomes for numerous rice cultivars. However, inferring gene functions, associated phenotypes, and causal variants from these sequences remains challenging. The Rice Annotation Project Database (RAP-DB; https://rapdb.dna.affrc.go.jp) is a curated genomic resource that provides comprehensive gene annotations for the reference genome of *Oryza sativa* ssp. *japonica* cv. ‘Nipponbare.’ Since its major update in 2013, gene models and functional annotations have been continuously revised through expert manual curation of newly published literature related to rice genes. As of March 2025, a total of 6,631 transcripts corresponding to 6,371 loci have been curated based on 4,699 peer-reviewed publications. These curated genes are functionally characterized and are frequently associated with agronomic traits, including yield components, stress tolerance, and disease resistance. To support molecular breeding, RAP-DB now provides a curated catalogue of 904 agronomically important loci, including gene symbols, functional descriptions, and associated traits, together with more than 1,000 functionally characterized alleles compiled from the literature. In addition to in-house expert curation, RAP-DB integrates community-curated datasets for major gene families, such as WRKY transcription factors, S-domain receptor-like kinases, and leucine-rich repeat-containing receptors, thereby expanding coverage of key regulatory and defense-related genes. RAP-DB also incorporates reanalyzed RNA sequencing expression profiles alongside microarray-based expression data and co-expression networks, offering gene-centric views of expression patterns across tissues, conditions, and developmental stages. Furthermore, RAP-DB is linked to genome-wide variation datasets from diverse rice varieties through the TASUKE+ genome browser, enabling exploration of allelic diversity across varieties. To enhance annotation quality and long-term sustainability, AI-assisted literature screening and a web-based feedback system have been introduced, allowing users to submit corrections to gene models and report newly characterized genes or relevant publications. Together, these developments strengthen RAP-DB as a primary, literature-based gene annotation resource and provide a practical foundation for molecular breeding in rice.

## Introduction

Rice (*Oryza sativa*) serves as a staple food for nearly half of the global population and plays a central role in global food security. The genome sequence of the *japonica* rice cultivar ‘Nipponbare’ was completed in 2004 by the International Rice Genome Sequencing Project (IRGSP) and has since been updated several times (International Rice Genome Sequencing Project 2005; Kawahara et al. 2013). Although high-quality genome assemblies, including telomere-to-telomere (T2T) assemblies of multiple rice cultivars, have recently been generated using advanced sequencing technologies such as PacBio, Oxford Nanopore Technologies (ONT), and Hi-C, the latest IRGSP Nipponbare reference genome (Os-Nipponbare-Reference-IRGSP-1.0) remains widely used as the primary reference for rice genomics (Wang et al. 2018; Qin et al. 2021; Zhang et al. 2022; Shang et al. 2023; Guo et al. 2025). In parallel, comprehensive gene annotations based on this reference genome have been developed and are publicly available through databases such as the Rice Annotation Project Database (RAP-DB), the Rice Genome Annotation Project (RGAP), and the Information Commons for Rice (IC4R) (Sakai et al. 2013; Sang et al. 2020; Hamilton et al. 2025). Together, these resources have provided foundational knowledge across diverse areas of rice biology, genetics, and breeding.

At the same time, our understanding of rice gene functions and their associated traits has expanded substantially, with a growing number of studies reporting functional characterization of individual genes (Fig. S1). This expansion accelerated following the release of the rice reference genome sequence and its gene annotation (International Rice Genome Sequencing Project 2005; Ohyanagi et al. 2006), which established a fundamental platform for functional genomics. Progress in this field has been driven in part by the accumulation of large-scale genomic and transcriptomic datasets enabled by rapid advances in sequencing technologies. In addition, the advent of genome editing technologies has facilitated precise functional validation of genes, further accelerating discoveries in rice functional genomics.

The resulting scientific outputs, including literature describing functional studies and high-throughput sequencing datasets, have been deposited in public repositories such as PubMed for literature and the International Nucleotide Sequence Database Collaboration (INSDC) databases (SRA/DRA/ENA) for sequencing data. To fully exploit these literature-based functional insights and large-scale genomic and transcriptomic datasets for functional genomics and molecular breeding, organizing and integrating them into genome annotation databases that provide efficient access, cross-referencing, and reuse for the research community is essential.

Over the past decade, rapid advances in functional genomics and breeding technologies, including marker-assisted selection and speed breeding, have made molecular breeding based on rational design a practical strategy for crop improvement. This approach relies on detailed knowledge of the genes underlying key agronomic traits (Zeng et al. 2017; Watson et al. 2018; Chukwu et al. 2019; Agata et al. 2020; Wei et al. 2021; Tian et al. 2021; Yang et al. 2023). Recent developments in genome editing technologies have enabled precise modifications not only of protein-coding regions but also of regulatory elements involved in transcription, splicing, and translation, resulting in phenotypic changes in traits such as flowering time, amylose content, and grain size (Sukegawa et al. 2022; Zhang et al. 2025). Furthermore, targeted modifications of transcriptional regulatory elements have been shown to fine-tune complex traits, including panicle branching and multi-pathogen resistance, by modulating gene expression levels (Kuroha et al. 2025; Han et al. 2025). These strategies ultimately depend on accurate genetic dissection of agronomic traits and high-resolution information on the underlying genes and alleles.

Several databases have been developed to compile information on rice gene functions and associated agronomic traits extracted from published literature. For example, funRiceGenes systematically curates gene function data from published studies, providing a comprehensive resource for exploring gene functions, phenotypes, and relevant literature (Huang et al. 2022). An additional challenge in rice genomics is inconsistent gene nomenclature, as individual genes are often referred to by multiple names across studies. Oryzabase addresses this issue by compiling gene names and symbols reported in the literature, thereby facilitating the resolution of nomenclature inconsistencies (Yamazaki et al. 2010). In addition to nomenclature, Oryzabase provides extensive information on mutants, quantitative trait loci (QTLs), genetic markers, and genomic diversity. Furthermore, Q-TARO/OGRO offers curated datasets of agronomically important rice QTLs and genes derived from the literature, contributing to the identification of trait-associated loci relevant to breeding programs (Yonemaru et al. 2010; Yamamoto et al. 2012). Comprehensive databases that integrate genomic and transcriptomic information with literature-derived functional annotations, therefore, represent valuable resources for advancing molecular breeding.

The gene annotation in the current version of RAP-DB was originally constructed for the IRGSP-1.0 genome in 2013 (Sakai et al. 2013). Gene models were generated based on alignments of full-length cDNA sequences from *Oryza sativa* ssp. *japonica*, mRNA sequences from monocot species (including rice, wheat, barley, and maize), and ab initio gene predictions, resulting in the prediction of 45,998 loci. Functional annotations were assigned using computational approaches, including sequence similarity searches against GenBank mRNA and UniProt protein databases, as well as functional domain prediction using InterProScan, which also provided Gene Ontology (GO) term assignments.

Since the major update in 2013, RAP-DB has been continuously maintained and enhanced through the incorporation of new datasets, additional functionalities, and improvements to the user interface. To further support the rice research community, expert-driven manual curation has been conducted to refine gene models and improve functional annotations. More recently, curated information on agronomically important genes and their functional alleles has been made available to facilitate molecular breeding research. RAP-DB also integrates transcriptome profiles from more than 700 RNA sequencing samples and genome-wide polymorphism data from approximately 700 diverse rice varieties. By combining literature-based curation with transcriptomic and genome diversity resources, RAP-DB continues to serve as a central platform for rice genomics and breeding research. This paper describes recent developments and ongoing activities in RAP-DB.

## Materials and Methods

### Manual Curation of Gene Models and Functional Annotations

Manual curation was conducted to update gene structures and functional annotations based on newly published literature. Relevant articles were identified by expert curators through keyword-based searches in PubMed (Sayers et al. 2021). For each curated gene, functional information, gene symbols, gene names, cellular localization, and related traits described in the literature were examined and used to revise existing RAP-DB annotations. When necessary, gene models were updated based on reported transcript structures, sequence information, and other supporting evidence. External resources, including gene annotations from RGAP (https://rice.uga.edu/) and RefSeq (https://www.ncbi.nlm.nih.gov/refseq/), as well as RNA sequencing and Iso-Seq read alignments, were also consulted to validate and refine the updated gene models. In addition, information on known allelic variants reported in the literature for genes associated with agronomically important traits was collected. This included details such as variant type and genomic position, the cultivars in which the alleles were identified, and their effects on phenotypic traits.

The process of updating gene models and functional annotations in RAP-DB is managed using an in-house web application developed specifically to support curation workflows. During each update, InterProScan is re-executed for all genes using the latest program version and database sources (Jones et al. 2014; Blum et al. 2025). For the latest annotation release, published on 19 March 2025, InterProScan v5.73-104 and the most recent version of the go.obo file downloaded from the Gene Ontology Resource website (https://geneontology.org/) were used to assign InterPro domains and Gene Ontology (GO) terms (Gene Ontology Consortium et al. 2023). In addition, gene-associated information from Oryzabase (https://shigen.nig.ac.jp/rice/oryzabase/), including gene symbols, gene names, Trait Ontology (TO), and Plant Ontology (PO) terms, was updated to reflect the latest available data. Enriched and depleted ontology terms were tested using Fisher’s exact test, and multiple testing correction was performed using the false discovery rate (FDR) method (Benjamini and Hochberg 1995).

To compare gene annotations across major rice annotation databases, protein sequence datasets were downloaded from RAP-DB (https://rapdb.dna.affrc.go.jp/download/irgsp1.html), RGAP (https://rice.uga.edu/download_osa1r7.shtml), RefSeq (https://www.ncbi.nlm.nih.gov/genome/annotation_euk/Oryza_sativa_Japonica_Group/102/), and IC4R (https://ngdc.cncb.ac.cn/ic4r/download). The completeness of each annotation set was evaluated using BUSCO v5.8.2 with the embryophyta_odb10 reference dataset (Manni et al. 2021).

### AI-Assisted Literature Classification for Manual Curation

To train the text classification model, a labeled dataset consisting of positive and negative literature sets was assembled. The positive dataset comprised 4,699 articles previously used for manual curation. For each publication, titles and abstracts were used as input data. The negative dataset consisted of an equal number of rice-related articles randomly selected from PubMed using the keywords “rice” and “Oryza sativa,” excluding articles already included in the positive dataset. When full-text content was available, it was used to extract additional information, such as gene identifiers and gene symbols, to support manual curation tasks.

To classify literature relevant to rice gene function, two natural language processing (NLP) models were implemented: fastText and BERT (Bojanowski et al. 2016; Joulin et al. 2016; Devlin et al. 2018). For fastText, the Python fasttext library v0.9.3 was used with default parameter settings. For BERT, the implementation provided in the transformers library v4.37.2 was employed. The final model was trained using the following hyperparameters: 50 epochs, a batch size of 16 for training and 32 for evaluation, 500 warm-up steps, a weight decay of 0.01, and a learning rate of 1 × 10⁻⁷. Five-fold cross-validation was performed using 80% of the labeled dataset for training and 20% for testing in each fold. Model performance was evaluated using precision, recall, and F1 score to assess effectiveness in selecting literature for manual curation.

### Transcriptome Sequencing and Data Analysis

For Iso-Seq analysis, rice plants (cv. Nipponbare) were grown in a growth chamber, and fully expanded leaves were sampled every two hours over a 24-hour period, one month after sowing. At each time point, one leaf was harvested from each of three individual plants, resulting in three leaves per time point. Total RNA was extracted from all samples using the Promega Maxwell 16 LEV simplyRNA Tissue Kit and then pooled in equal amounts to obtain an RNA sample for Iso-Seq library preparation. Iso-Seq libraries were constructed using a PacBio Iso-Seq kit, and three size-selected libraries (1–2 kb, 2–3 kb, and 3–6 kb) were prepared. Sequencing was performed on the PacBio RS II platform using two SMRT cells for the 1–2 kb library, three cells for the 2–3 kb library, and four cells for the 3–6 kb library. The resulting reads were processed using the Iso-Seq analysis pipeline v3.0.1, which included extraction of full-length non-chimeric reads, clustering of similar transcripts, and polishing to generate high-quality consensus sequences (Guizard et al. 2023). Polished transcripts were mapped to the reference genome (Os-Nipponbare-Reference-IRGSP-1.0) (Kawahara et al. 2013), and redundant isoforms were removed based on the alignments using cDNA_cupcake v16.0.0 (https://github.com/Magdoll/cDNA_Cupcake). In total, 24,172 non-redundant Iso-Seq transcripts were obtained. Complete coding sequences (CDSs) were predicted for each transcript using TransDecoder v5.5.0 (https://github.com/TransDecoder/TransDecoder).

RNA sequencing data for 746 publicly available Nipponbare samples representing diverse tissues and growth conditions were obtained from public databases (SRA/DRA/ENA) (Table S1). Raw sequencing data were preprocessed using Trimmomatic v0.39 with the parameters: ILLUMINACLIP:illumina_adapter_sequences.fa:2:30:10, LEADING:15, TRAILING:15, SLIDINGWINDOW:10:15, and MINLEN:30 (Bolger et al. 2014). Clean reads were aligned to the reference genome using HISAT2 v2.2.1 with the parameters --min-intronlen 20 and --max-intronlen 10000 (Kim et al. 2019). Transcript abundances were estimated as transcripts per kilobase million (TPM) using StringTie v2.2.1 (Pertea et al. 2015).

### Detection of genome-wide variations

Publicly available whole-genome resequencing data from diverse rice varieties were obtained from public databases (SRA/DRA/ENA) (Table S2). Raw reads were preprocessed using Trimmomatic v0.38 with the parameters: ILLUMINACLIP:adapters.fa:2:30:10, LEADING:20, TRAILING:20, SLIDINGWINDOW:10:20, and MINLEN:30. Filtered paired-end reads were aligned to the reference genome using BWA-MEM v0.7.17 (Li and Durbin 2009). PCR duplicates were identified and removed using Picard v2.18.17 (https://broadinstitute.github.io/picard/), and variant calling was performed using the HaplotypeCaller module in GATK v4.2.4.0 (Van der Auwera and O’Connor 2020). Joint genotyping across all samples was conducted using the GenotypeGVCFs module in GATK.

Following genotyping, variant filtering was applied to obtain high-confidence single-nucleotide polymorphisms (SNPs) and insertions/deletions (InDels). Variants with sequencing depth outside the range of the average depth ± five standard deviations were excluded. For SNPs, additional filtering criteria included QD < 2.0, MQ < 40.0, FS > 60.0, SOR > 3.0, MQRankSum < −12.5, and ReadPosRankSum < −8.0. For InDels, the criteria were QD < 2.0, FS > 200.0, SOR > 10.0, and ReadPosRankSum < −20.0. Filtered variants were annotated using SnpEff v5.2f (Cingolani et al. 2012) to predict potential effects on gene structure and function based on the latest RAP-DB gene annotation. The annotated variant dataset was implemented in TASUKE+ version 20241122 together with the latest RAP-DB gene annotation (Kumagai et al. 2019).

Biallelic SNPs were extracted from the genome-wide variation dataset using PLINK v1.9 (Chang et al. 2015). SNPs with a minor allele frequency (MAF) < 0.05 or a missing genotype rate > 10% were excluded. The resulting high-confidence SNP dataset was used as input for ADMIXTURE v1.3.0 (Alexander et al. 2009), which was run with values of K (number of ancestral populations) ranging from 5 to 8.

## Results

### Refinement of Rice Gene Annotations through Continuous Literature-based Manual Curation

Since the major update in 2013 (Sakai et al. 2013), RAP-DB has been continuously updated with revised gene models and functional annotations, with new releases issued approximately twice per year. The latest RAP-DB gene annotation release, dated 19 March 2025, comprises 45,870 loci and 53,144 transcript isoforms (Table 1). To enhance functional characterization, InterProScan has been applied to all annotated transcripts at each release to assign protein functional domains and Gene Ontology (GO) terms. In the most recent release, functional domains were detected in 26,815 loci (58.5%), and 18,946 loci (41.3%) were annotated with at least one GO term.

**Table 1.**
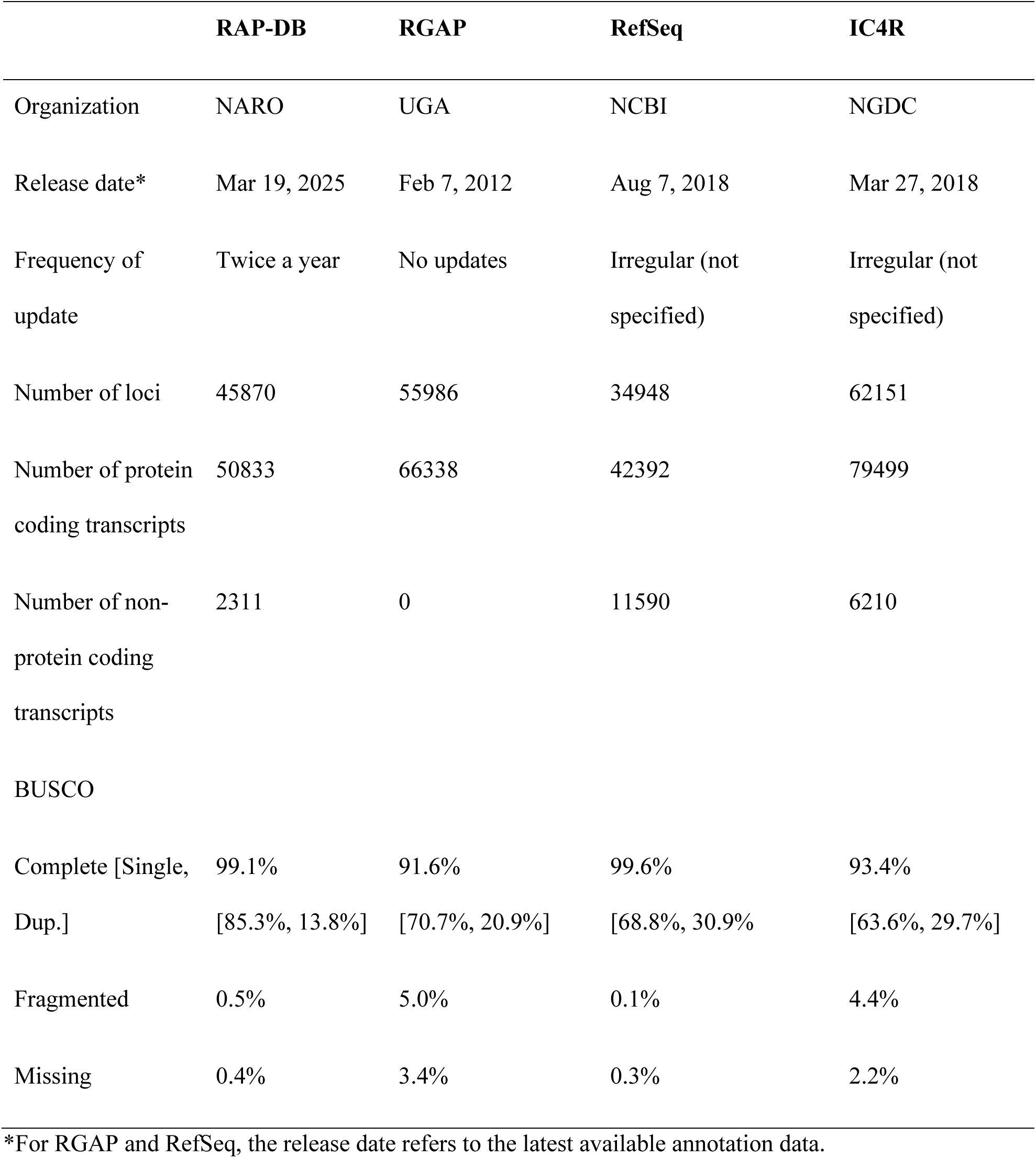
Overview of rice gene annotation databases.

|  | **RAP-DB** | **RGAP** | **RefSeq** | **IC4R** |
| --- | --- | --- | --- | --- |
| Organization | NARO | UGA | NCBI | NGDC |
| Release date* | Mar 19, 2025 | Feb 7, 2012 | Aug 7, 2018 | Mar 27, 2018 |
| Frequency of update | Twice a year | No updates | Irregular (not specified) | Irregular (not specified) |
| Number of loci | 45870 | 55986 | 34948 | 62151 |
| Number of protein coding transcripts | 50833 | 66338 | 42392 | 79499 |
| Number of non-protein coding transcripts | 2311 | 0 | 11590 | 6210 |
| BUSCO |  |  |  |  |
| Complete [Single, Dup.] | 99.1%<br>[85.3%, 13.8%] | 91.6%<br>[70.7%, 20.9%] | 99.6%<br>[68.8%, 30.9%] | 93.4%<br>[63.6%, 29.7%] |
| Fragmented | 0.5% | 5.0% | 0.1% | 4.4% |
| Missing | 0.4% | 3.4% | 0.3% | 2.2% |

\*For RGAP and RefSeq, the release date refers to the latest available annotation data.

Through collaboration with Oryzabase (Yamazaki et al. 2010), manually curated rice gene names, symbols, and associated Trait Ontology (TO) and Plant Ontology (PO) terms have been incorporated into RAP-DB. Consequently, 6,598 loci (14.4%) were annotated with TO terms, 3,996 loci (8.7%) were associated with PO terms, and 14,983 loci (32.7%) were assigned gene names and symbols curated in Oryzabase.

To date, manual curation based on newly published literature has refined the gene models and functional annotations of 6,631 transcripts corresponding to 6,371 loci (13.9%), drawing on evidence from 4,699 peer-reviewed publications (Fig. 1). These curated entries are clearly indicated in RAP-DB by a “Curated” label, signifying that their gene models and functional annotations have been reviewed and refined by expert curators (Fig. 2A).

**Fig. 1.**
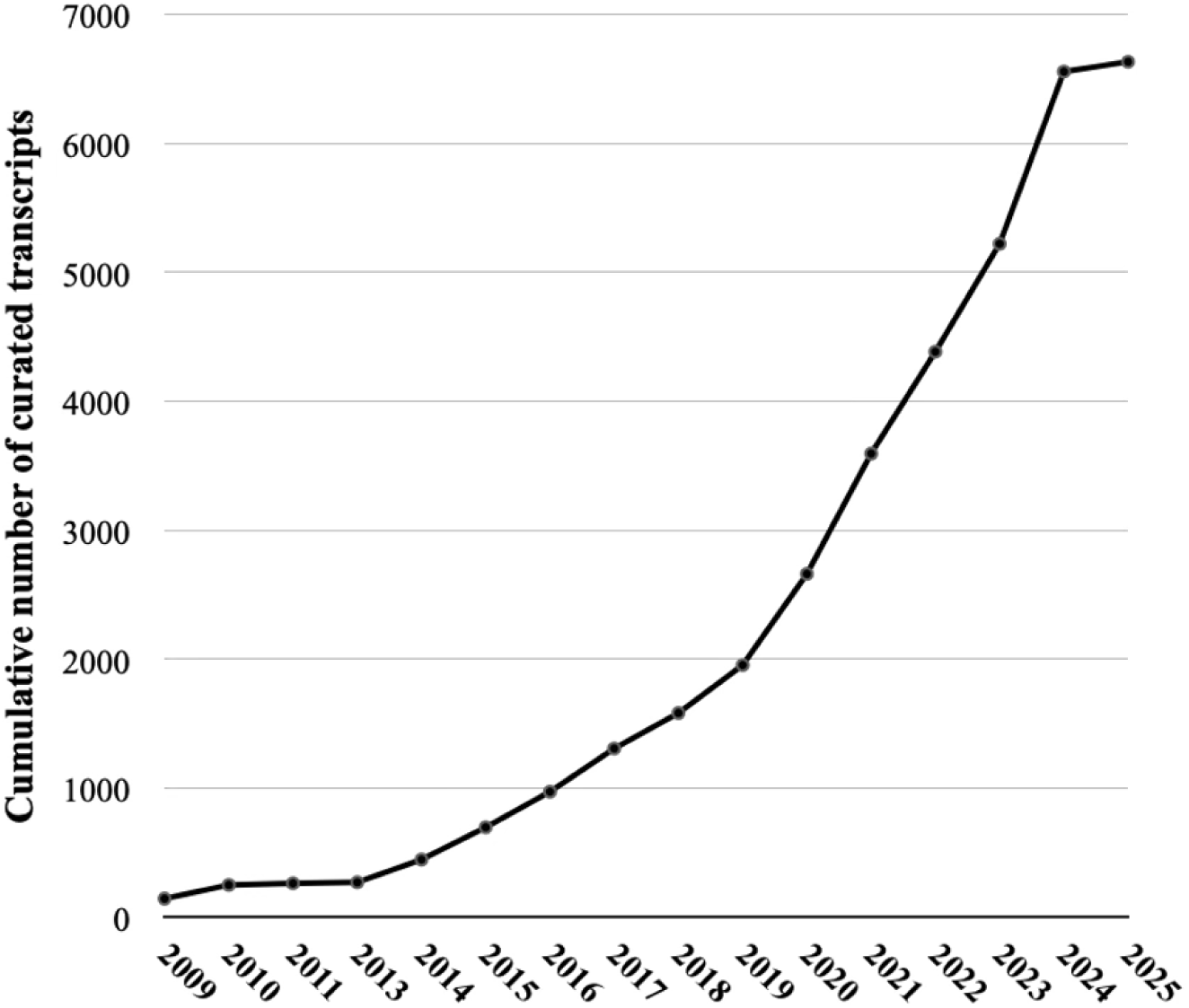
Trends in the number of manually curated genes in RAP-DB. Annual increase in the number of curated transcripts in RAP-DB from 2013 to March 2025. As of March 2025, 6,631 transcripts corresponding to 6,371 loci have been manually curated based on peer-reviewed publications.

**Fig. 2.**
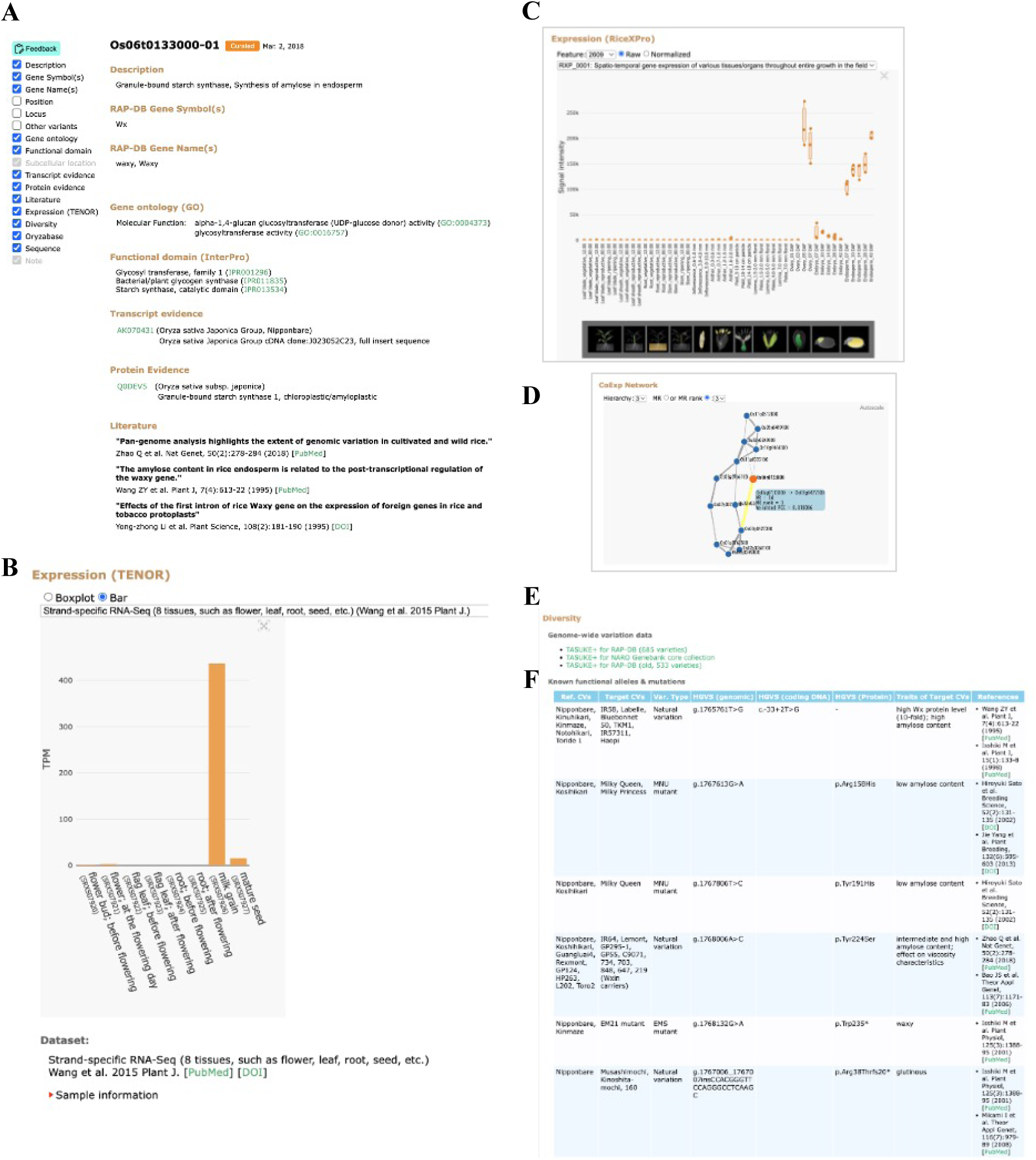
Annotation features of the transcript/locus information page in RAP-DB. (A) Curated annotation features in a transcript information page that are updated through manual curation, as indicated by the “Curated” label with the date of curation. Curated features include description of gene products and functions, gene symbols and names, transcript and protein evidence, and supporting peer-reviewed publications. Gene ontology (GO) terms and functional domains (InterPro) are updated through re-analysis at each update cycle to reflect the latest information. (B) Gene expression profiles across 746 RNA-Seq samples derived from 45 datasets, along with metadata for each sample. (C) A direct link to TASUKE+, enabling users to explore local variant data around the gene region. (D) Known functional alleles and mutations extracted from peer-reviewed publications. (E) Microarray-based gene expression profiles previously provided by RiceXPro. (F) Gene co-expression network information previously provided by RiceFREND.

Manual curation involves not only the extraction of gene functions and associated phenotypes but also the verification and, where necessary, revision of gene structures using multiple lines of supporting evidence. As RAP-DB serves as one of the primary annotation databases for the rice reference genome and provides official rice gene identifiers (e.g., Os06g133000 or Os06t133000-01) (McCouch and CGSNL, 2008), it plays a central role in maintaining consistent and accurate gene models. This framework enables the correction of misannotated gene structures and the incorporation of newly identified genes when supported by reliable evidence.

To ensure accurate gene structures, RAP-DB integrates information reported in the literature with annotations from other major resources, including RGAP (Hamilton et al. 2025) and RefSeq. In addition, RNA-Seq and Iso-Seq transcript alignments are used as transcriptomic evidence to support gene model validation. These datasets are visualized in JBrowse, allowing direct comparison with RAP-DB gene models (Fig. S2).

As a result of these sustained efforts, the quality of gene annotation has improved substantially compared with the computationally predicted gene models released in 2013. The BUSCO completeness score for the curated gene models increased to 99.1%, compared with 79.4% for the original annotation, demonstrating the effectiveness of continuous manual curation in improving annotation quality and completeness (Table S3).

To further characterize manually curated genes, we examined the distribution of TO, PO, GO, and InterPro domain terms among curated loci (Tables S4–S9). TO analysis revealed enrichment of traits related to environmental stress tolerance, including salt (TO:0006001), drought (TO:0000276), cold (TO:0000303), and heat tolerance (TO:0000259), as well as yield-related traits such as plant height (TO:0000207), tiller number (TO:0000346), seed set percentage (TO:0000455), and grain- and panicle-related attributes. Genes involved in disease resistance, including blast disease (TO:0000074) and bacterial blight resistance (TO:0000175), were also frequently represented (Fig. 3; Table S8).

**Fig. 3.**
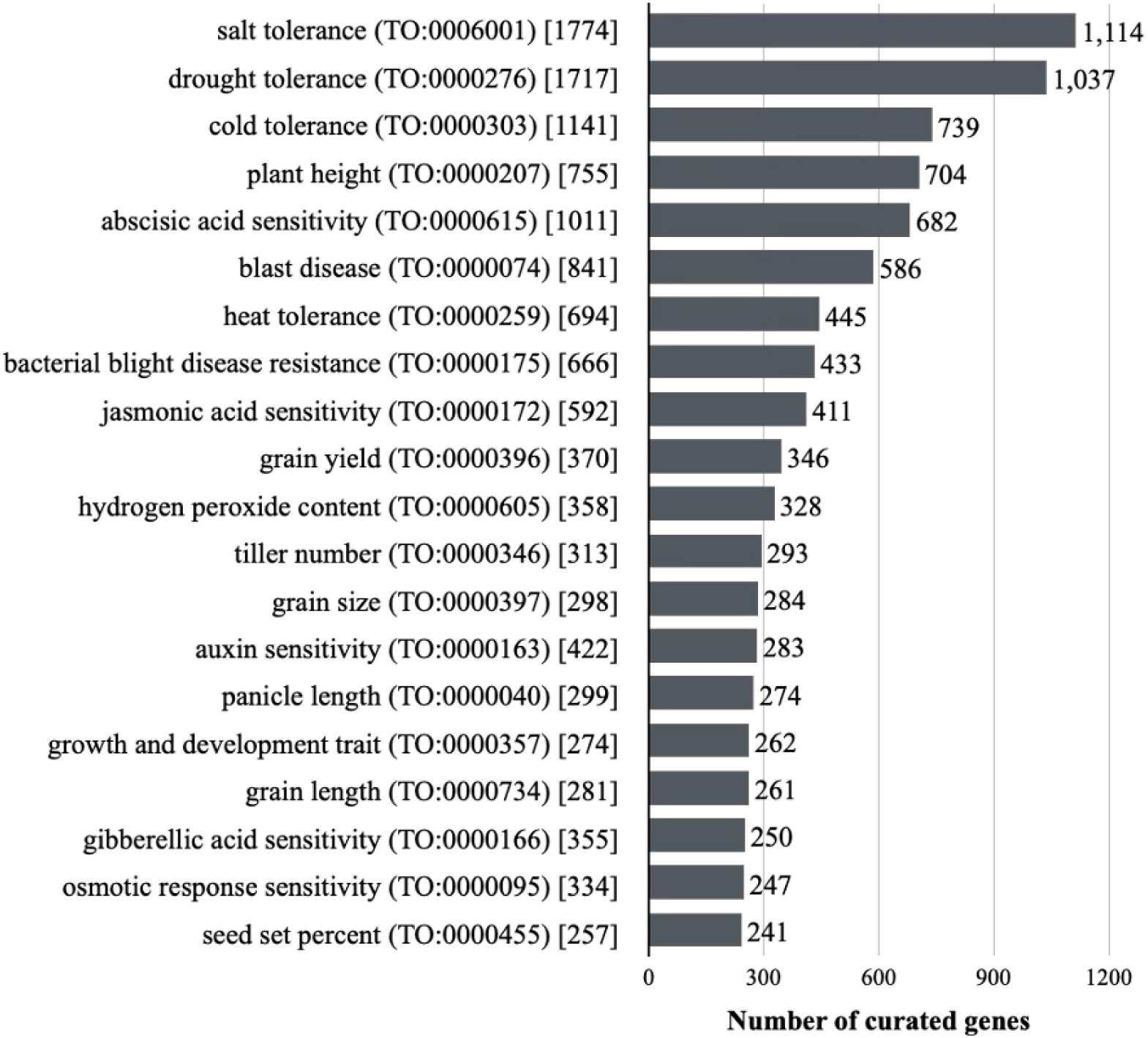
Top 20 Trait Ontology (TO) terms significantly enriched in the curated genes. The number of curated genes assigned to each of the top 20 enriched TO terms. The numbers in square brackets following each TO term indicates the total number of genes associated with the term. All TO terms are significantly over-represented in the curated genes (FDR < 0.01).

PO term analysis showed significant enrichment for terms associated with leaf (PO:0025034), root (PO:0009005), and inflorescence (PO:0009049), indicating that these organs and tissues are frequent targets of functional studies. GO term analysis revealed that curated genes were commonly annotated with terms such as ATP binding (GO:0005524), protein binding (GO:0005515), protein kinase activity (GO:0004672), protein phosphorylation (GO:0006468), and regulation of DNA-templated transcription (GO:0006355) (Fig. S3). These terms are typically linked to regulatory processes, including signal transduction and transcriptional control, suggesting that a large proportion of curated genes encode regulatory proteins. In contrast, GO terms associated with housekeeping functions, such as structural constituent of ribosome (GO:0003735), translation (GO:0006412), RNA modification (GO:0009451), and ribosome (GO:0005840), were underrepresented among curated genes.

InterPro domain analysis revealed significant enrichment of domains associated with disease resistance, including the leucine-rich repeat domain superfamily (IPR032675), leucine-rich repeat (IPR001611), and NB-ARC (IPR002182) domains. Domains linked to regulatory functions, such as the protein kinase-like domain superfamily (IPR011009), winged helix-like DNA-binding domain superfamily (IPR036388), and P-loop–containing nucleoside triphosphate hydrolase (IPR027417), were also enriched (Fig. S4). By contrast, domains such as the alpha/beta hydrolase fold (IPR029058) and F-box domain (IPR001810), which are typically associated with metabolic enzymes and ubiquitin-mediated proteolysis, respectively, were less frequent among curated genes.

### Enhancing Manual Curation with AI-Assisted Literature Selection

To facilitate the identification of publications suitable for functional annotation of rice genes, we developed an AI-assisted system to evaluate newly published literature for relevance to manual curation. Two machine-learning models, fastText (Joulin et al. 2017; Bojanowski et al. 2017), and Bidirectional Encoder Representations from Transformers (BERT) (Devlin et al. 2018), were evaluated for this task. fastText is a lightweight word-embedding classifier optimized for speed and accuracy, whereas BERT is a deep contextual language model that captures bidirectional relationships in text.

Using five-fold cross-validation, both models achieved F1 scores close to 0.9, indicating high predictive performance (Fig. S5A). To assess generalizability, the trained models were applied to an independent dataset of 216 newly published articles, and predictions were compared with curator assessments. Both models achieved F1 scores of approximately 0.8 in this external evaluation, demonstrating robust performance on previously unseen data (Fig. S5B). In addition, articles judged suitable for curation by curators consistently received higher relevance scores from both models, indicating their utility in supporting literature selection (Fig. S5C).

To integrate this functionality into routine curation workflows, we developed an in-house, web-based application termed “AI Curator” (Fig. S6). This tool retrieves articles from PubMed based on user-defined keywords and publication dates and automatically evaluates their relevance using the trained AI model. In addition to article titles and relevance scores, the application displays abstracts and, when available, gene identifiers and gene symbols extracted from full-text content. Incorporation of AI Curator into the curation workflow has substantially improved efficiency by enabling curators to prioritize high-confidence publications and accelerate functional annotation in RAP-DB.

### Community-Driven Curation to Sustain and Enrich RAP-DB Annotations

While expert manual curation has been central to ensuring annotation accuracy and reliability, this approach is inherently time-consuming and labor-intensive. In recent years, increasing contributions from the rice research community have enabled more efficient and comprehensive updates to gene annotations.

A representative example is the work of the Rice WRKY Working Group, which reconciled inconsistent nomenclature for WRKY transcription factors across publications (Rice WRKY Working Group 2012). WRKY genes form a large transcription factor family involved in diverse processes, including biotic and abiotic stress responses, senescence, and development. Using this curated dataset, we revised the gene symbols and names of 75 WRKY loci in RAP-DB to reflect unified nomenclature. In addition, two recently curated gene family resources were integrated into RAP-DB. Naithani et al. (2021) cataloged 144 members of the S-domain subfamily of receptor-like kinases (SDRLKs), which participate in signaling pathways related to development and stress responses. Gottin et al. (2021) reported more than 1,000 leucine-rich repeat–containing receptor (LRR-CR) genes, many of which are receptor-like kinases involved in pathogen recognition and plant innate immunity. These datasets were used to update gene identifiers, gene symbols, and names, and functional descriptions for the corresponding RAP-DB entries.

To further promote community participation, we recently introduced a user feedback system that allows researchers to propose gene model revisions and submit information on newly characterized genes or relevant publications (Fig. 2A; Fig. S7). This initiative supports collaborative curation and helps ensure that RAP-DB remains current and accurate.

These community-driven updates, together with literature-based expert curation, likely contribute to the sustained growth in RAP-DB usage. In 2024, RAP-DB received an average of 680,000 page views and 37,400 visits per month (Fig. S8A). Although most users access RAP-DB from Japan and China, substantial usage also originates from the United States, India, South Korea, and Taiwan, underscoring its importance to the global rice research community (Fig. S8B).

### Comprehensive Gene Expression Profiles and Co-expression Networks

As a satellite database of RAP-DB, TENOR was previously released to provide gene expression profiles for leaves and roots under various abiotic stress conditions (Kawahara et al. 2016). Since then, numerous transcriptome studies conducted under diverse cultivation conditions and in mutant backgrounds have been deposited in public repositories. We retrieved RNA sequencing datasets comprising 746 Nipponbare samples spanning a wide range of tissues and cultivation conditions, including those previously included in TENOR, and reanalyzed them using a unified pipeline based on the latest RAP-DB gene models (Table S1).

Because metadata associated with RNA sequencing datasets in INSDC repositories are often incomplete or inconsistent, we manually curated sample attributes, such as tissue or organ, cultivation conditions, and library construction methods, from the original publications. To standardize descriptions across studies, Plant Ontology (PO) and Plant Experimental Conditions Ontology (PECO) terms were assigned to each sample. The resulting expression profiles are displayed on each transcript information page as part of the functional annotation (Fig. 2B), and read alignment tracks for all RNA sequencing samples are available in the genome browser (Fig. S2). These alignments also serve as transcriptional evidence for validating and refining gene models.

In parallel, RiceXPro and RiceFREND have long served as community resources providing large-scale microarray-based expression profiles and co-expression networks across tissues and developmental stages (Sato et al. 2013a, b). RAP-DB integrates RiceXPro expression profiles and RiceFREND co-expression networks with curated gene annotations and reanalyzed RNA sequencing data, presenting them together on each locus page (Fig. 2C–D). Collectively, these datasets indicate where genes are expressed and with which partners they are co-expressed, providing a valuable resource for functional inference and prioritization of candidate genes for downstream analyses.

### Variant Landscapes Across Diverse Rice Varieties in TASUKE+

To support interpretation of phenotypic diversity and discovery of alleles of interest, we compiled whole-genome resequencing datasets from diverse rice varieties and identified SNPs and indels using a unified analysis pipeline. The resulting variation data are provided through TASUKE+, an interactive genome browser that enables visualization of genome-wide polymorphisms across multiple varieties (Kumagai et al. 2019).

RAP-DB currently hosts three major variation datasets through TASUKE+: (i) 685 rice varieties collected from multiple published studies (Table S9); (ii) the NARO Genebank Core Collection, comprising 119 accessions from the World Rice Core Collection and the Japanese Rice Core Collection(Tanaka et al. 2020, 2021), and (iii) the NARO Genebank Open Rice Collection, consisting of 623 accessions (Tanaka et al. 2025). TASUKE+ is tightly integrated with RAP-DB annotations, allowing users to open genomic regions directly from transcript information pages to visualize local variation, compare allelic states among varieties, and inspect nucleotide-level differences (Fig. 2E; Fig. S9). TASUKE+ also provides tools for constructing phylogenetic trees based on user-defined genomic regions and for designing PCR primers around target variants, facilitating downstream experimental validation and molecular breeding.

For the 685-variety dataset, resequencing reads were retrieved from public repositories (SRA/DRA/ENA) based on 44 publications (Table S2). Variety-level metadata, including names, subgroups, and geographic origins, were manually curated from the literature and database records. This collection includes both cultivated rice and wild *Oryza* accessions, capturing broad genomic diversity among AA-genome *Oryza* species and supporting studies of evolution, domestication, and adaptation (Fig. S10).

### A Curated Catalogue of Agronomically Important Genes and Functional Alleles

Although TASUKE+ enables genome-wide visualization of polymorphisms, the phenotypic consequences of individual variants are not immediately apparent. To bridge genotype and phenotype for molecular breeding, we compiled literature-derived information on agronomically important genes and their functional alleles and linked curated annotations to specific variants.

In the current release, RAP-DB provides a curated list of 904 agronomically important loci (“Agri. genes”) associated with traits such as flowering time, yield components, grain quality, stress tolerance, and disease resistance (Fig. 4). For each locus, the list includes the gene symbol or name, associated traits (with TO terms when available), and relevant literature references. In addition, functionally characterized alleles reported in peer-reviewed publications were manually curated and linked to the corresponding transcript pages (Fig. 2F). Each allele entry summarizes the variant type (e.g., SNP or indel), the cultivars carrying the allele, and the reported phenotypic effects, together with the source reference.

**Fig. 4.**
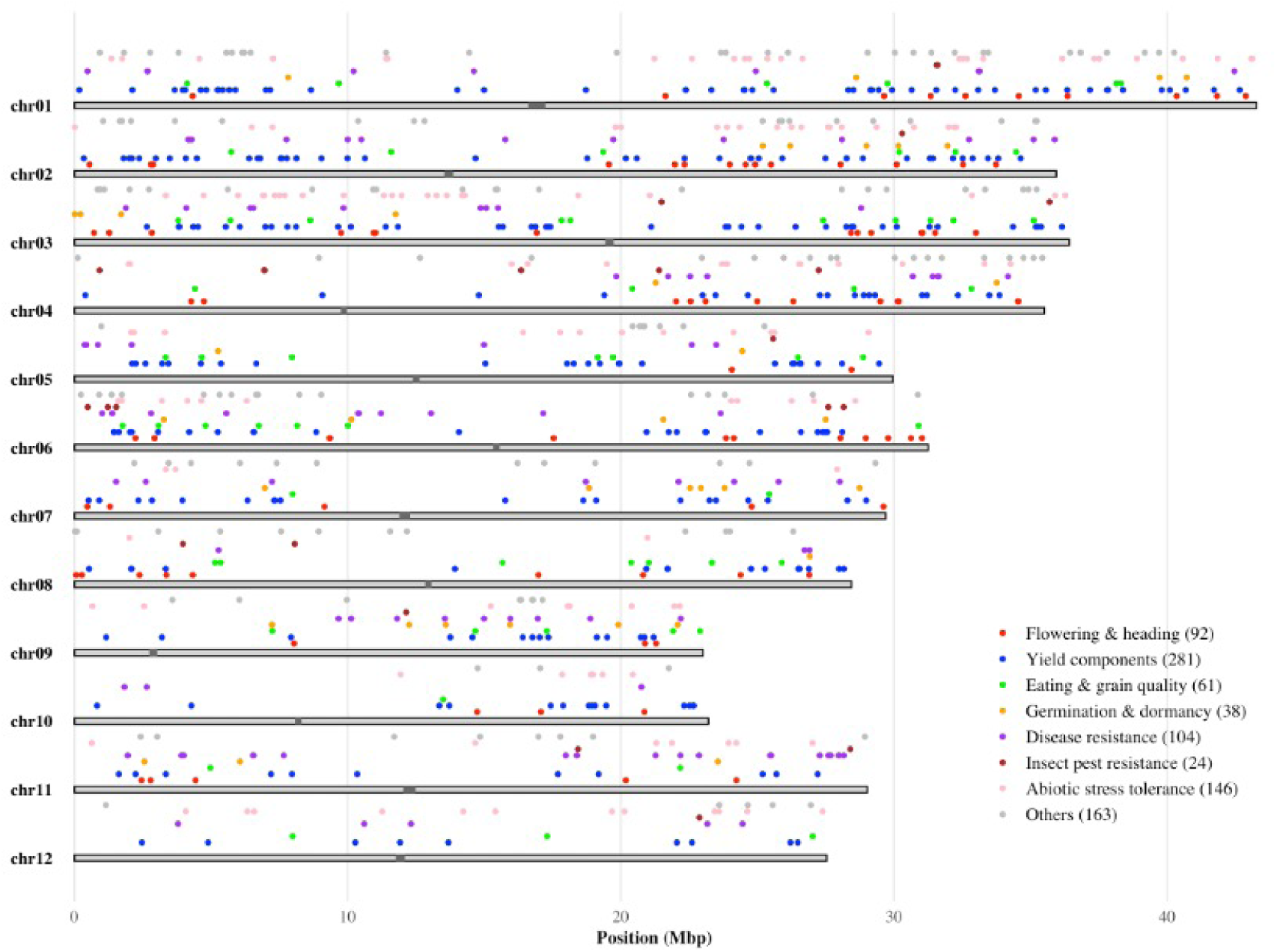
Chromosomal distribution of curated agronomically important genes (Agri. genes). Agri. genes are plotted on the 12 chromosomes and color-coded by trait category. The number of genes in each category is indicated in parentheses. Trait categories include: flowering and heading, yield components, eating and grain quality, germination and dormancy, disease resistance, insect pest resistance, abiotic stress tolerance, and others.

The Agri. gene list supports filtering by gene symbol or name, trait keywords, and TO identifiers, enabling rapid identification of loci relevant to specific phenotypes (Fig. S11). For example, a search for “amylose content” retrieves 25 loci, including *Wx1* (Os06t0133000-01), which has 10 functionally characterized alleles associated with glutinous and high-amylose phenotypes listed on its transcript information page (Fig. 2F). Together with TASUKE+, these curated functional allele annotations provide a practical resource for identifying candidate causal variants and supporting downstream analyses in breeding workflows.

## Discussion

Recent advances in sequencing technologies have made it increasingly feasible to generate high-quality reference genome assemblies. Computational annotation pipelines enable rapid, genome-wide gene prediction and functional inference based on sequence homology to known proteins and functional domains. However, computational approaches still have limitations in accurately predicting gene function and cannot readily incorporate the latest findings reported in the literature. To address these limitations, RAP-DB has been actively maintained through expert, literature-based curation over the past 15 years, with regular biannual releases. As of 19 March 2025, 6,371 loci have been manually curated from 4,699 publications, yielding high-confidence functional annotations that reflect current knowledge of rice genes.

While funRiceGenes cites over 7,900 publications and Oryzabase indexes more than 26,000 articles related to rice genes, RAP-DB currently includes a larger number of functionally characterized genes than the approximately 4,500 genes reported by funRiceGenes. Unlike databases that primarily aggregate literature information, RAP-DB functions as a primary annotation database, in which curated findings are directly integrated into genome annotations by revising gene models and assigning updated gene identifiers. To our knowledge, this combination of continuous literature-driven updates and the authority to modify primary gene models is unique among rice gene–related databases.

Traditionally, the identification of studies relevant to rice gene function relied on curators manually searching PubMed (Sayers et al. 2021). However, as the volume of rice-related literature has continued to expand, this approach has become increasingly time-consuming and inefficient. Recent advances in artificial intelligence (AI), particularly in natural language processing (NLP), have substantially improved document classification accuracy using machine-learning approaches. In this study, we implemented AI-assisted literature selection to improve the efficiency of manual curation. In parallel, we increasingly incorporate curated gene family datasets reported in peer-reviewed studies, including those for the WRKY, SDRLK, and LRR-CR gene families. RAP-DB also collaborates closely with Oryzabase, regularly exchanging gene models, symbols, names, and publication information to improve both resources. To further broaden participation, RAP-DB now provides a user feedback system that allows researchers to suggest gene model revisions and submit newly published evidence of gene function. All external contributions are reviewed using standardized procedures to ensure annotation accuracy and consistency prior to integration.

RAP-DB will continue to provide up-to-date rice gene annotations by integrating expert literature curation, AI-assisted literature screening, and carefully reviewed community input, while directly updating primary gene models when new, reliable evidence becomes available. Large volumes of next-generation sequencing data are deposited in public repositories such as SRA, DRA, and ENA, yet many datasets are used only once in their original studies and are rarely reused. In addition, heterogeneous bioinformatics pipelines across publications hinder direct comparison of reported polymorphisms and expression values. To address these challenges, we reanalyze published whole-genome resequencing and RNA-Seq datasets using a standardized analysis pipeline and disseminate the results through RAP-DB and TASUKE+. This uniform processing framework enables direct cross-study comparisons across cultivars and experimental conditions.

Transcriptome data provide valuable evidence for gene existence and functional inference. RAP-DB integrates RNA-Seq datasets derived from 746 experimental conditions retrieved from public repositories and reanalyzed under a unified pipeline. In addition, we incorporate established transcriptome resources from RiceXPro, which provides microarray-based expression profiles, and RiceFREND, which offers co-expression networks based on RiceXPro data. Together, these resources provide a comprehensive view of gene activity across tissues, developmental stages, and experimental conditions, and are displayed alongside curated annotations within RAP-DB.

To support reproducibility, we provide full documentation of the tools and parameters used in our analysis pipelines, enabling researchers to reproduce our results or process their own datasets in a compatible manner. In parallel, we curated and normalized sample metadata, including tissue or organ, cultivation conditions, variety names, and subgroup information, using controlled vocabularies such as Plant Ontology (PO) and Plant Experimental Conditions Ontology (PECO).

We will continue to expand RAP-DB by incorporating additional genome assemblies from diverse rice varieties and wild relatives, as well as transcriptome datasets sampled across broader developmental stages and environmental conditions. These efforts will further strengthen RAP-DB as a standardized and comparable multi-omics resource for the rice research community. Early applications of DNA marker-based breeding focused on relatively simple strategies, such as introgression of a single resistance gene into elite genetic backgrounds. In contrast, modern breeding increasingly combines multiple loci that influence flowering time, yield components, grain quality, stress tolerance, and disease resistance(Zeng et al. 2017; Wei et al. 2021). At the same time, advances in genome editing technologies now enable precise manipulation of gene function, including targeted changes in coding regions and regulatory elements that affect transcription, splicing, and translation(Sukegawa et al. 2022).

In this context, comprehensive information on genes underlying agronomic traits, as well as the allelic variants present at those loci, is essential. Through manual literature-based curation, we extract detailed information on mutation types and phenotypic effects and integrate this knowledge into RAP-DB gene annotations. These curated gene and allele datasets are intended to support the design of elite cultivars, from selecting optimal allele combinations to identifying targets for genome editing.

Looking ahead, we will continue to expand both the breadth and granularity of RAP-DB resources. We will maintain ongoing literature-based curation of agronomically and biologically important genes and their alleles, while extending genome-wide functional variant annotation to improve interpretation of both coding and regulatory polymorphisms. Together, these efforts aim to provide a practical data foundation for precision breeding strategies that fully leverage genomic and gene-level knowledge.

## Conclusion

We report recent advances in RAP-DB, including long-standing efforts to manually curate gene structures and functions, while improving the efficiency and sustainability of these workflows; the integration of comprehensive transcriptome and genome-wide variation datasets; and the compilation of curated lists of agronomically important trait-associated genes along with their known functional alleles. Through continued expert curation and collaboration with the rice research community, RAP-DB now provides a rich, literature-supported repository of functionally characterized genes, and updates will continue to incorporate new findings promptly.

To further enhance both efficiency and coverage of curation, we are developing advanced AI-based approaches, including large language models capable of reading full-text articles to extract gene functions, associated traits, and causal variants, thereby accelerating the curation workflow. RAP-DB is poised to serve as a standard curated gene set for pan-genome studies across rice cultivars and *Oryza* species. Future developments will extend annotations by linking SNPs, indels, and structural variations across diverse cultivars and wild relatives. In parallel, we are constructing orthology mappings between rice genes and those of other crops, enabling cross-species transfer of knowledge and supporting comparative genomics and translational crop research.

## Abbreviations

CDS: coding sequence
FDR: False discovery rate
GO: Gene Ontology
IRGSP: International Rice Genome Sequencing Project
LRR-CR: Leucine-rich repeat-containing receptor
MAF: Minor allele frequency
NLP: Natural language processing
PECO: Plant Experimental Conditions Ontology
PO: Plant Ontology
RAP-DB: Rice Annotation Project Database
RGAP: Rice Genome Annotation Project
SDRLK: S-domain subfamily of receptor-like kinase
TENOR: Transcriptome Encyclopedia of Rice
TO: Trait Ontology
TPM: Transcripts Per Kilobase Million

## Declarations

### Ethics Approval and Consent to Participate

Not applicable

### Consent for Publication

Not applicable

### Availability of Data and Materials

The Iso-Seq sequence data analyzed in this study is available in the SRA/DRA/ENA under the following accessions: PRJDB8306.

RAP-DB https://rapdb.dna.affrc.go.jp

### Authors’ Contributions

Y.K., H.S. and T.I. conceived and designed the project. Y.K., H.S. supervised and coordinated the manual curation of gene annotations. T.H.K. and X.W. conducted manual curation of gene annotation and functional alleles. R.H. developed and maintained the database and managed the web server infrastructure. Y.T. developed the AI curator system. M.K. and N.T. analyzed genome diversity data and implemented new features of TASUKE+ for RAP-DB. Y.K. wrote the main manuscript. All co-authors reviewed and approved the final version.

## Acknowledgements

We would like to thank Kazuhiko Sugimoto, Akira Takahashi, Kiyosumi Hori, Hironori Itoh, Yasumori Tamura, Ken Ishimaru, Ritsuko Mizobuchi (NARO), and Mitsuhiro Obara (JIRCAS) for sharing their knowledge and suggesting functionally characterized genes related to agronomic traits. We would like to thank Shoko Kawamoto and Yo Shidahara (National Institute of Genetics, Oryzabase) for sharing gene symbols and names information, and for their valuable discussions. We would like to thank Toshiyuki Sato (Mizuho Research & Technologies) for valuable comments on the construction of AI models. We also thank the Research Center for Advanced Analysis, NARO for the use of the high-performance cluster computing system and web servers. A part of the computational analyses in this study were conducted by the NARO AI research supercomputer “Shiho”.

