## Supplemental Material 1 for "Rice Annotation Project Database (RAP-DB): literature-curated gene annotation and integrated omics resources for rice functional genomics and molecular breeding"

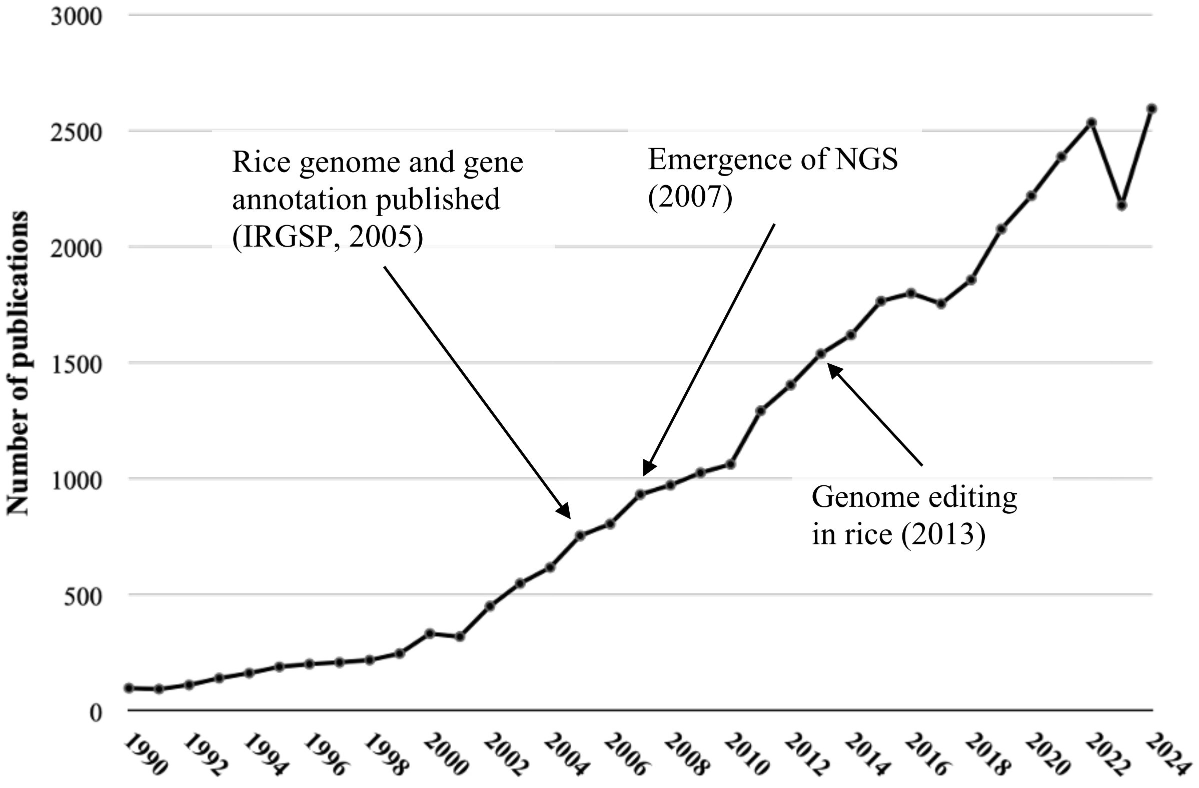


**Fig. S1** Annual number of rice-related publications retrieved from PubMed from January 1, 1990 to December 31, 2024. The number of publications per year was retrieved using the search query ("Oryza sativa" OR "rice") AND "gene" on PubMed. Arrows indicate three key technological milestones in rice functional genomics.

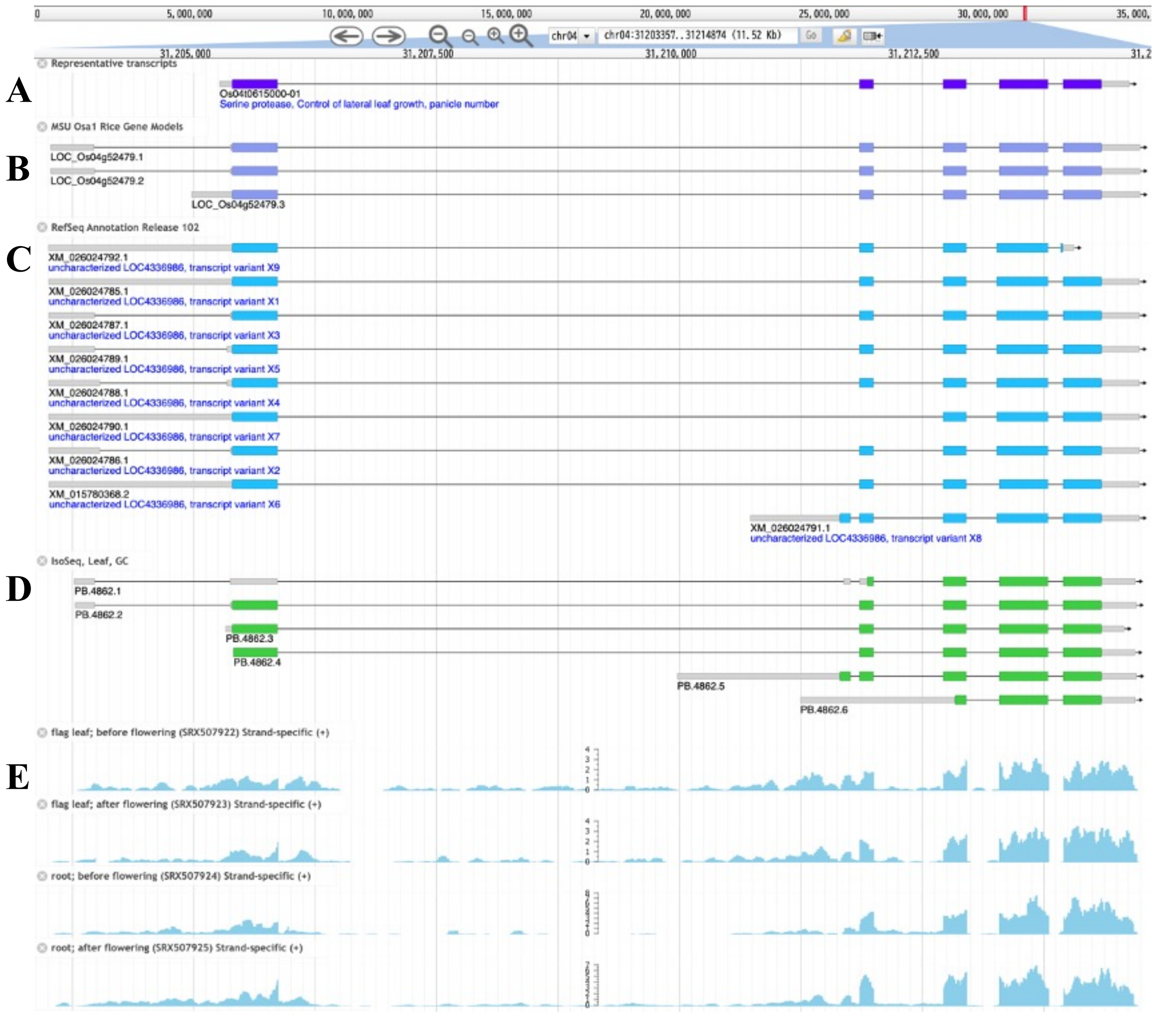


**Fig. S2** Visualization of gene models and transcriptional evidence for NAL1 in the RAP-DB JBrowse. The screenshot includes (A) the RAP-DB curated gene model, along with the gene model from (B) RGAP and (C) RefSeq. It also shows transcriptional evidence from (D) Iso-Seq and (E) RNA-Seq read alignments.

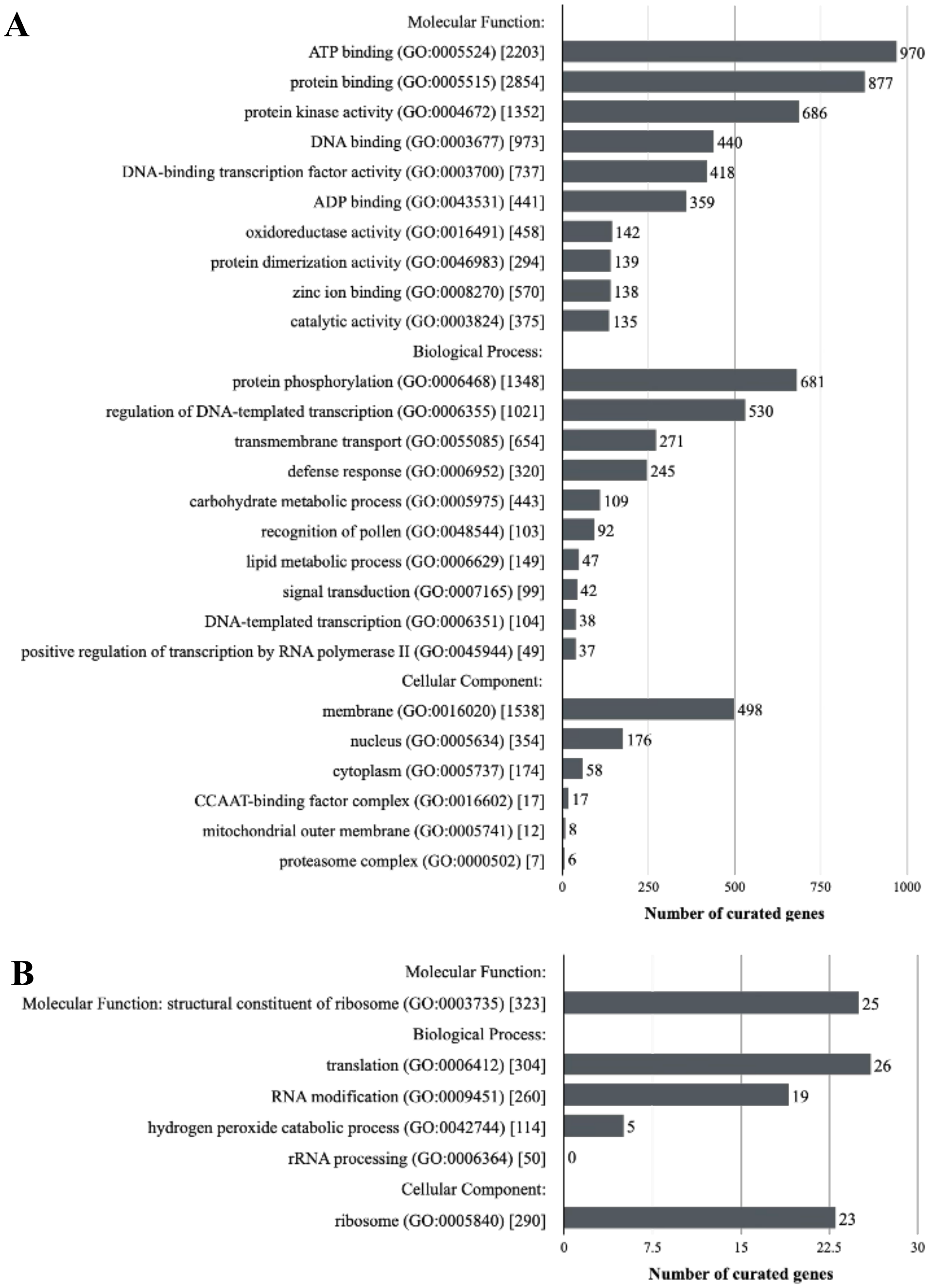


**Fig. S3** Gene Ontology (GO) terms that are significantly (FDR < 0.01) enriched (A) and depleted (B) among the curated genes compared to the whole gene set. Terms are grouped by GO category. The numbers in square brackets indicate the total number of genes annotated with each term in the whole gene set.

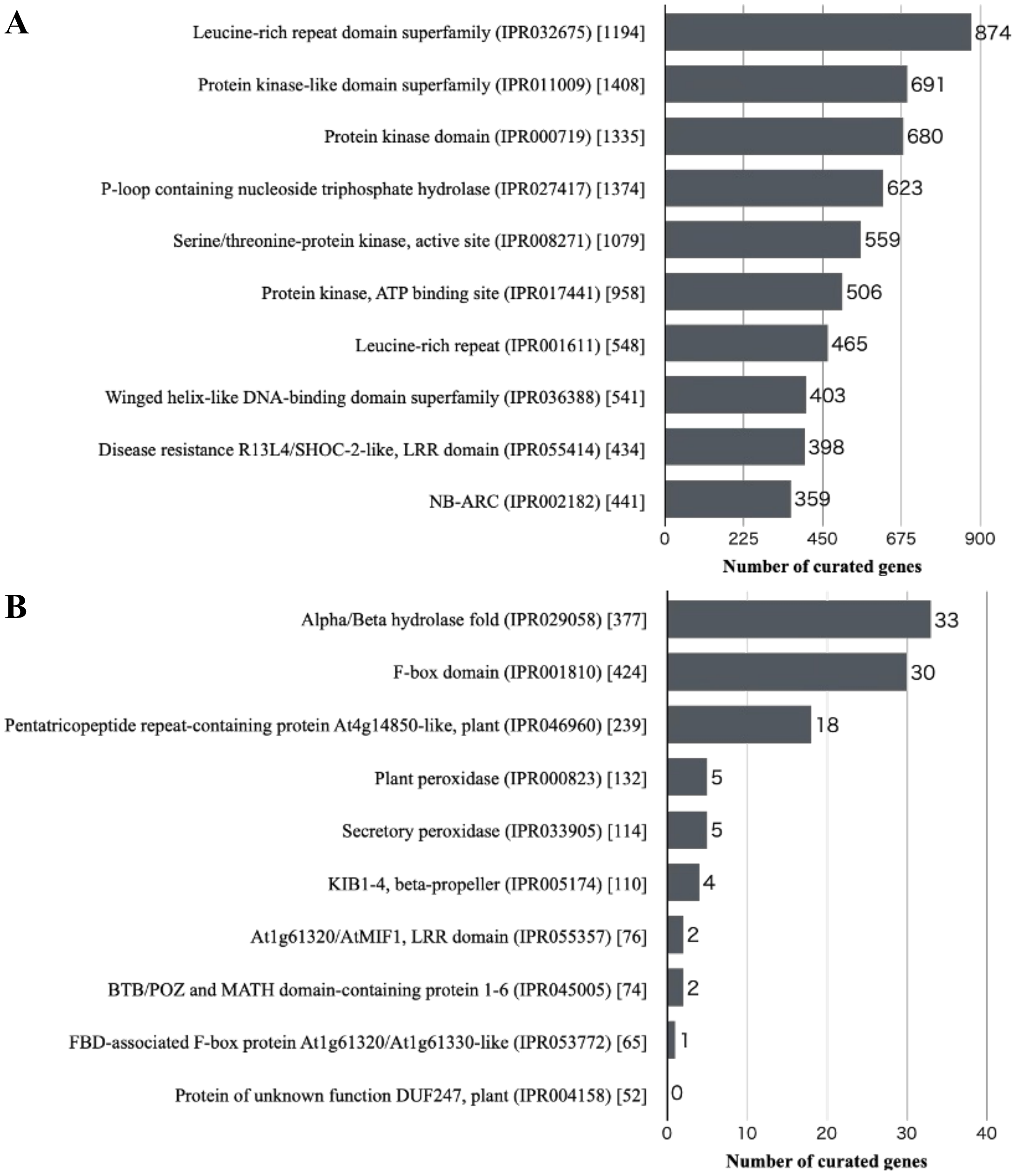


**Fig. S4** InterPro domains that are significantly (FDR < 0.01) enriched (A) and depleted (B) among the curated genes compared to the whole gene set. The numbers in square brackets indicate the total number of genes annotated with each domain in the whole gene set.

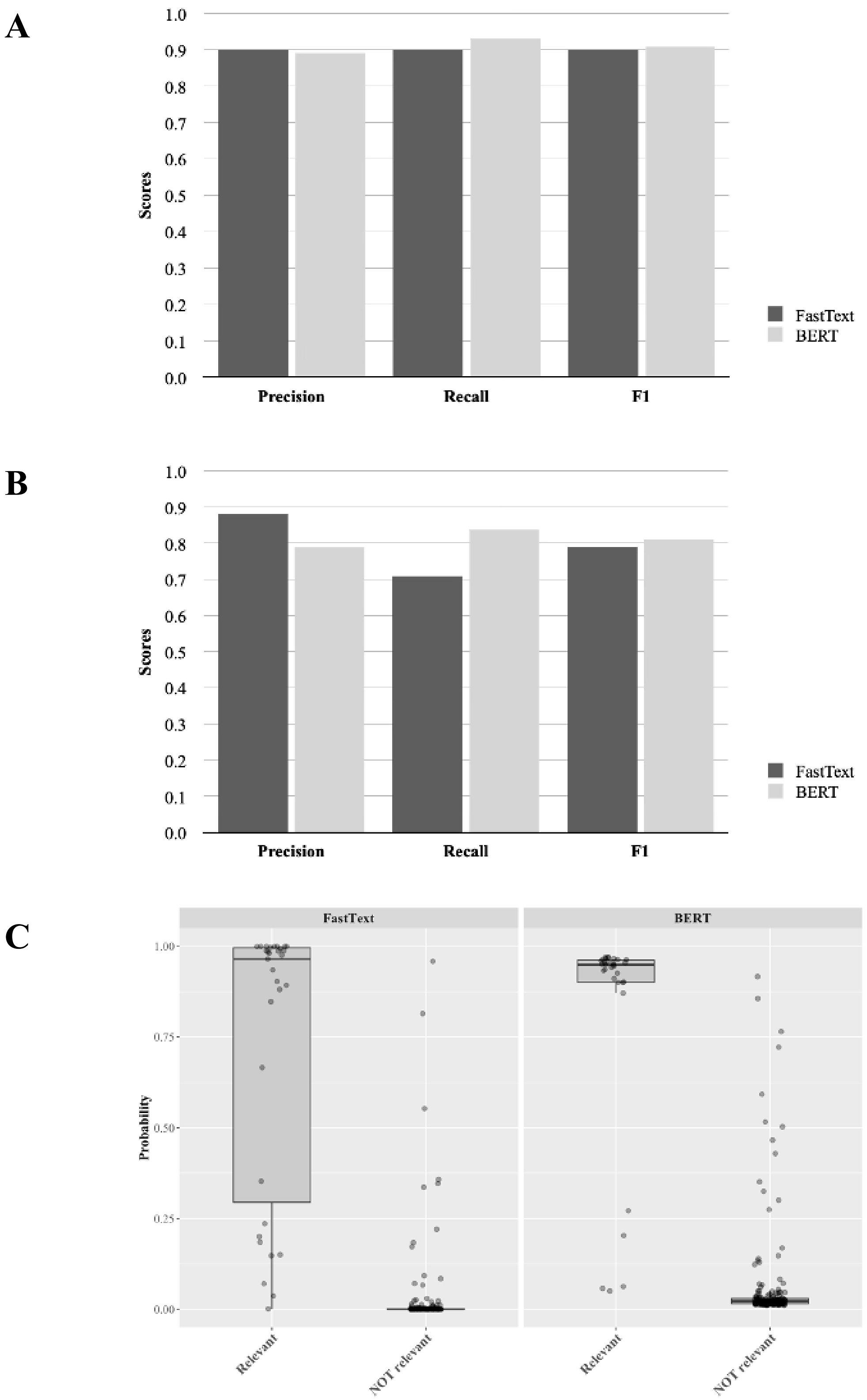


**Fig. S5** Performance evalution of fastText and BERT models for literature selection. Precision, recall, and F1 scores from (A) five-fold cross-validation and (B) external evaluation on 216 newly published articles. (C) Prediction probability distributions for articles judged as relevant or not by the curator.

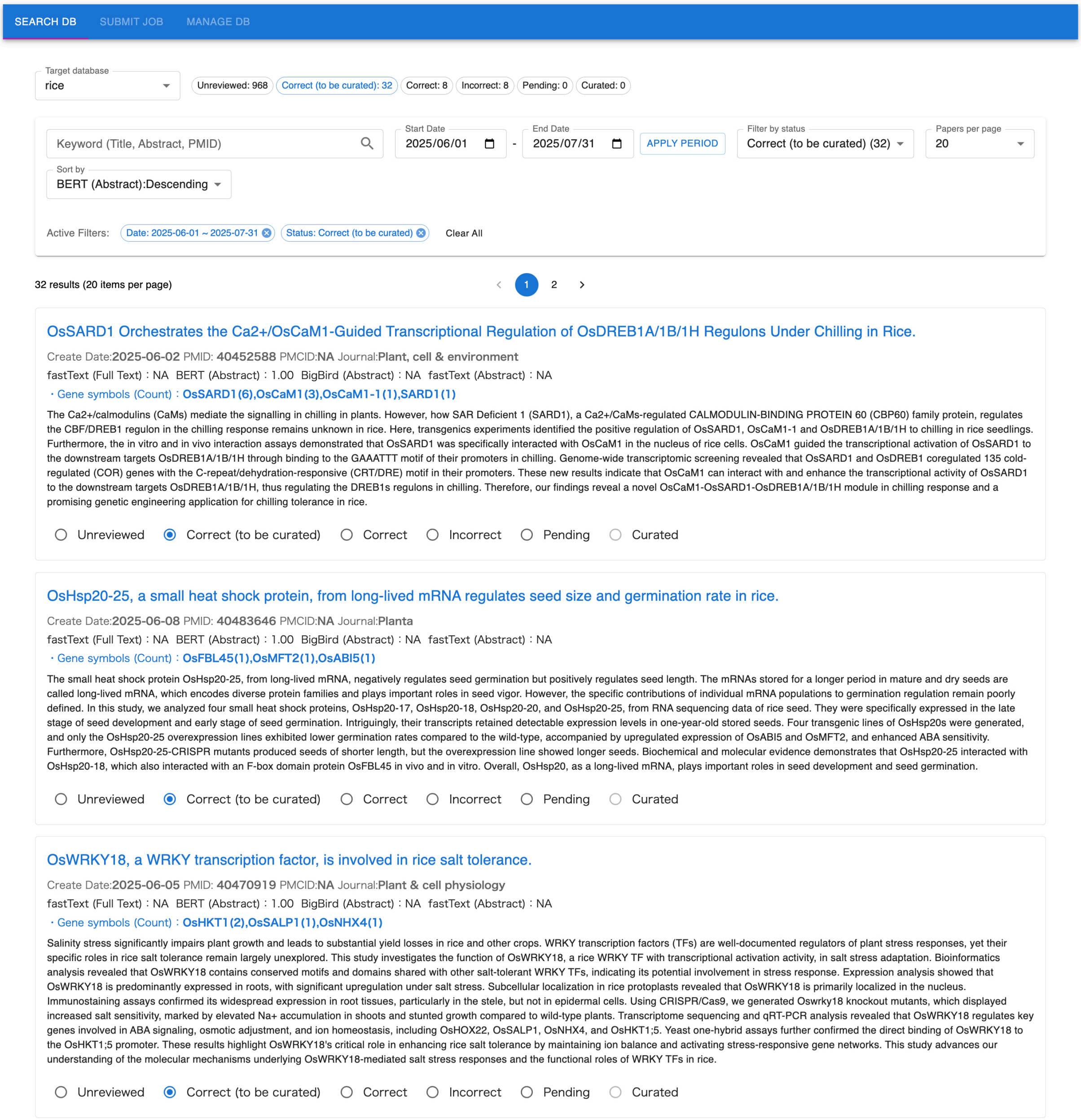


**Fig. S6** Screenshot of the in-house web-based application "AI Curator." The system retrieves literature from PubMed based on user-specified keywords and publication dates, and for each article displays the title, abstract, predicted relevance probability from the trained AI models, and, when available, gene IDs and gene symbols automatically extracted from the full text.

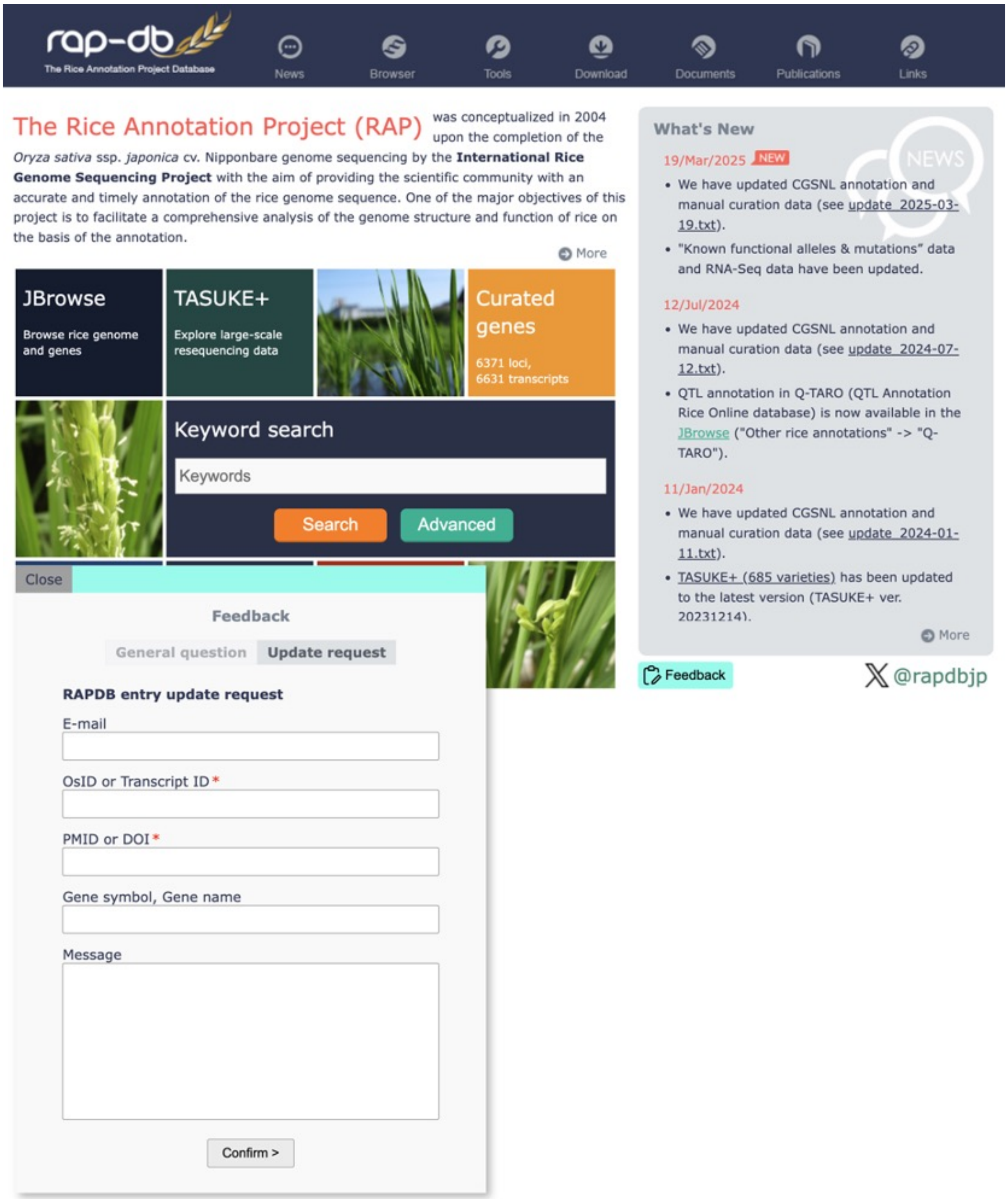


**Fig. S7** User feedback system interface in RAP-DB. The feedback system implemented in RAP-DB allows users to contribute to annotation improvements by submitting suggestions for gene model revisions and newly published functional information. Users can provide gene IDs, gene symbols and names, relevant publication references, and detailed comments.

**A**

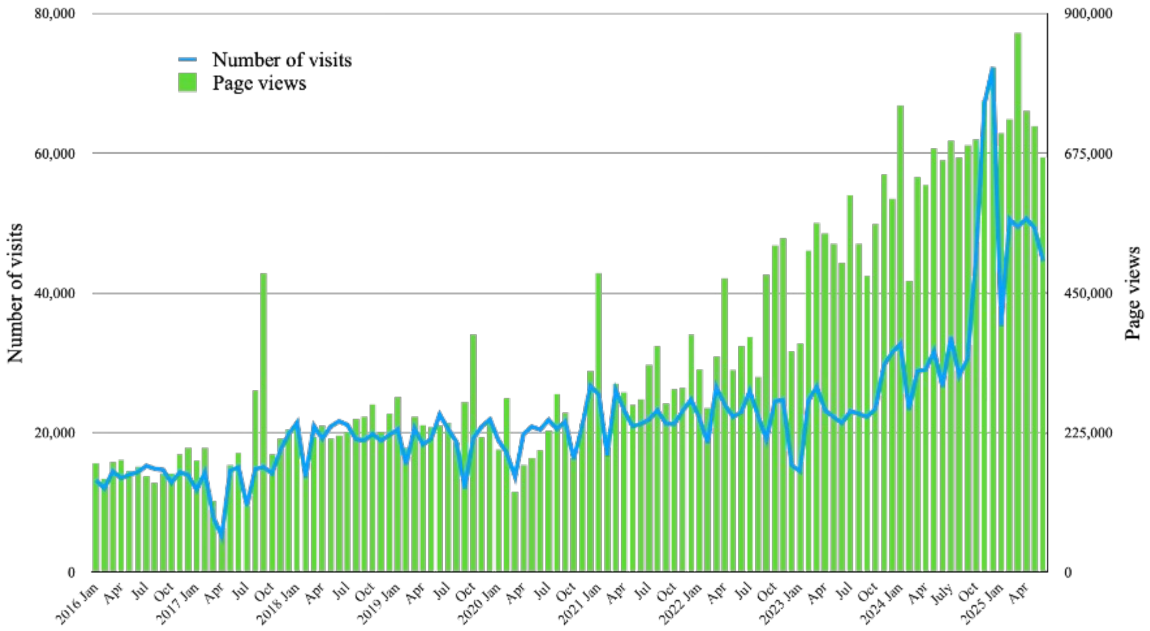


**B**

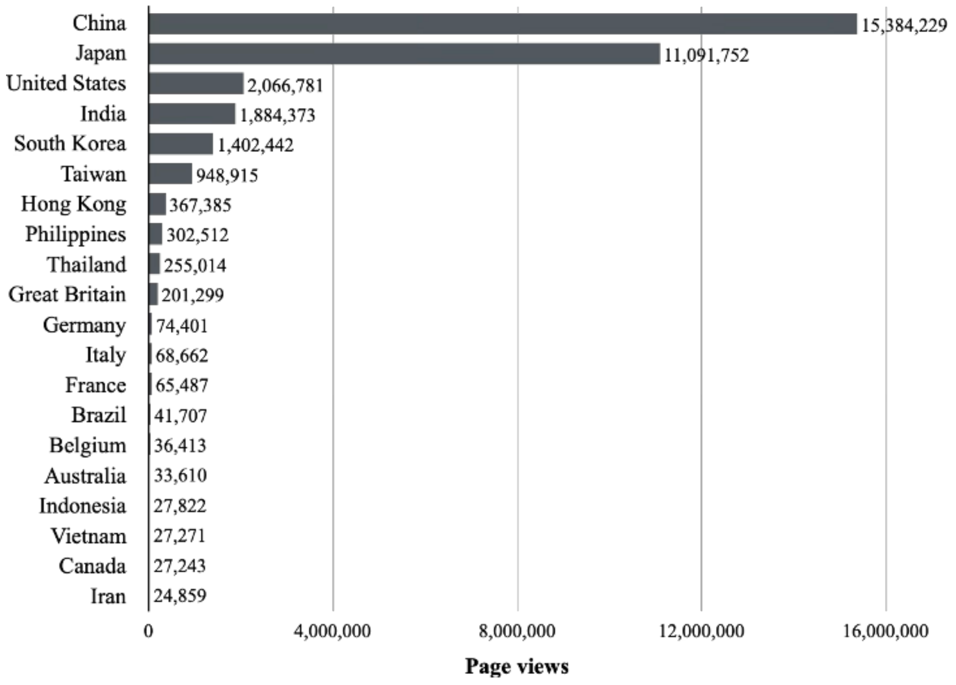


**Fig. S8** Access statistics of RAP-DB from January 1, 2016 to June 30, 2025. (A) Monthly access trends. The bar graph shows the number of page views per month, and the line graph indicates the number of visits. B) Total page views by country.

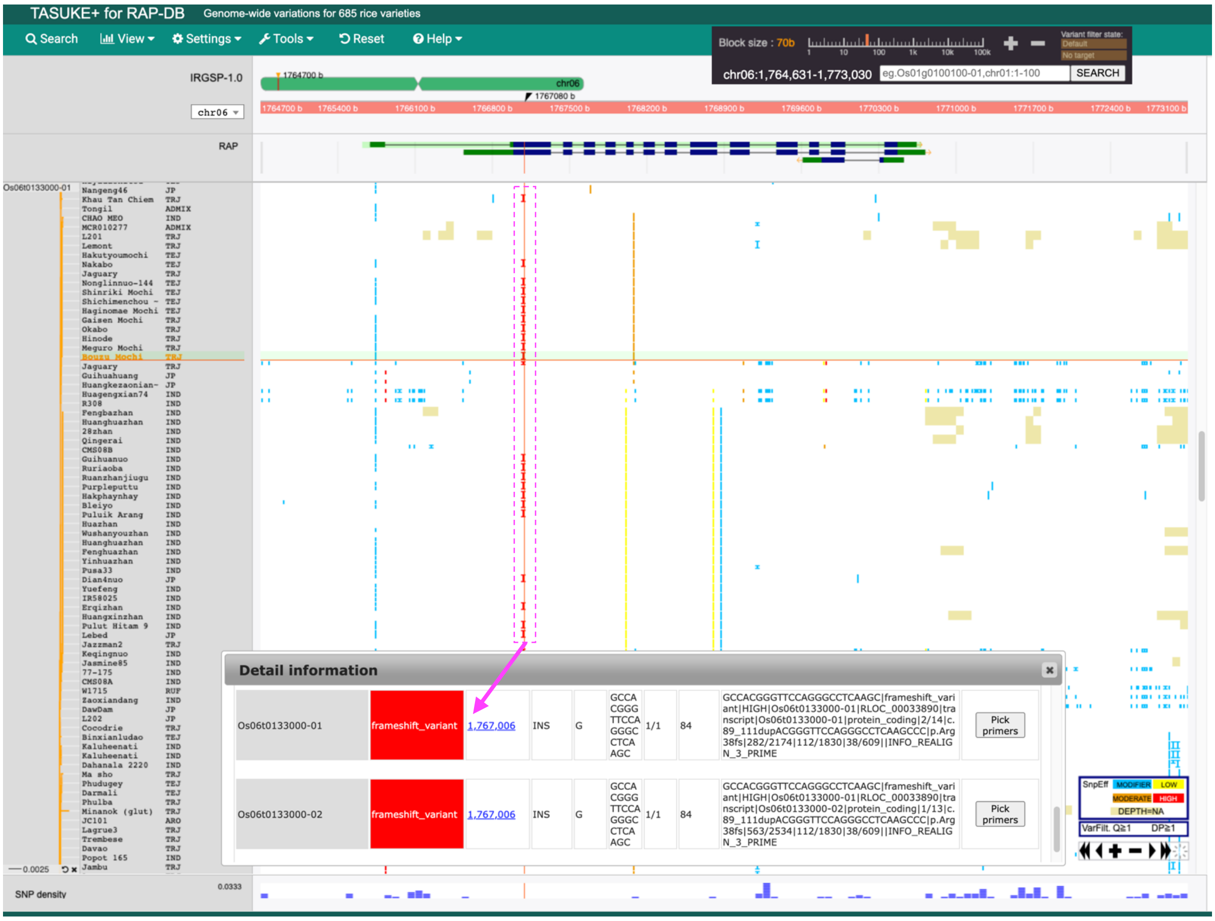


**Fig. S9** Screenshot of TASUKE+ highlighting the 24-bp insertion in the Wx locus, which has been reported in many glutinous japonica rice varieties (indicated by the dotted magenta rectangle). By clicking the 24-bp insertion in the main view, the detailed annotation for this variant is displayed in a pop-up window.

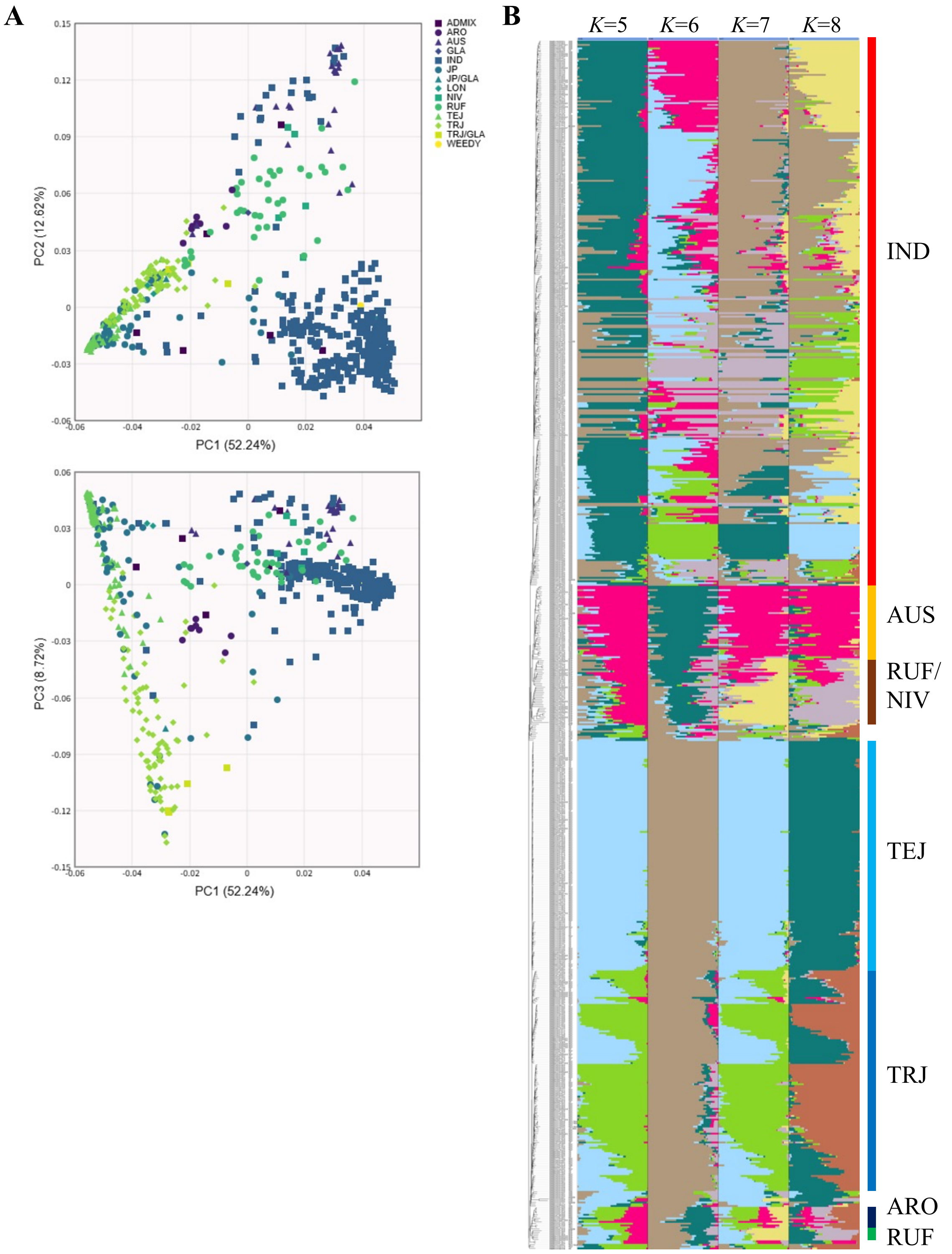


**Fig. S10** Population structure of the 685 cultivated and wild rice accessions with variant data available in TASUKE+. (A) PCA plots showing PC1 versus PC2 and PC1 versus PC3, calculated from LD-pruned biallelic SNPs with MAF > 0.05 across all accessions. (B) Population structure of all accessions inferred by ADMIXTURE with K = 5–8 ancestral clusters. Accessions are ordered according to the neighbor-joining tree shown on the left. Major varietal groups and species are indicated on the right. Abbreviations: RUF, *Oryza rufipogon*; NIV, *O. nivara*; IND, indica; AUS, aus; TEJ, temperate japonica; TRJ, tropical japonica; ARO, aromatic.

**Agronomically important genes**

amylose content | << | < | > | >> | Per page 10 | 11 - 20 of 26 | The number of agri. genes: **25** loci, **26** transcripts

| Locus ID | Transcript ID | Position | Gene symbols | Gene names | Trait Ontology | Plant Ontology | References | Related traits (in Japanese) |
|---|---|---|---|---|---|---|---|---|
| Os04g0413500 | Os04t0413500-01 | chr04:20422171..20426921 | GIF1, gif1, CIN2, OsCIN2, OsGIF1, WB1, OsWB1, GIF1/OsCIN2 | GRAIN INCOMPLETE FILLING 1, grain incomplete filling 1, "Beta-fructofuranosidase, insoluble isoenzyme 2", Sucrose hydrolase 2, Invertase 2, Cell wall beta-fructosidase 2, cell-wall invertase 2, White Belly 1 | chalky endosperm (TO:0000266)<br>filled grain number (TO:0000447)<br>grain size (TO:0000397)<br>amylopectin content (TO:0000097)<br>endosperm color (TO:0000487)<br>glume color (TO:0000221)<br>endosperm related trait (TO:0000575)<br>1000-dehulled grain weight (TO:0000592)<br>invertase activity (TO:0000311)<br>spikelet number (TO:0000456)<br>amylose content (TO:0000196)<br>seed maturation (TO:0002661)<br>sucrose content (TO:0000328)<br>fructose content (TO:0006005)<br>seed size (TO:0000391)<br>grain weight (TO:0000590)<br>grain yield (TO:0000396)<br>seed quality (TO:0000162)<br>starch content (TO:0000696)<br>seed thickness (TO:0000304)<br>seed width (TO:0000149)<br>seed length (TO:0000146)<br>starch grain shape (TO:0002656)<br>seed set percent (TO:0000455)<br>glucose content (TO:0000300) | inflorescence internode (PO:0006326)<br>endosperm development stage (PO:0007633)<br>endosperm (PO:0009089)<br>pericarp (PO:0009084)<br>root tip (PO:0000025)<br>carpel vascular system (PO:0005019) | Cho JI et al. Plant Cell Rep, 24(4):225-36 (2005) [PubMed]<br>Wang E et al. Nat Genet, 40(11):1370-4 (2008) [PubMed]<br>Wang H et al. Int J Mol Sci, 19(8): (2018) [PubMed] | 食味・品質 |
| Os02g0725900 | Os02t0725900-01 | chr02:30191485..30192488 | HAP3K, OsHAP3K, OsHAP3K/OsNF-YB1, OsNF-YB1, NF-YB1, nf-yb1, OsLEC1, OsNF-YB-1, NFYB1, OsEnS-41 | HAP3K SUBUNIT OF CCAAT-BOX BINDING COMPLEX, Nuclear transcription factor Y subunit B-1, CCAAT-binding transcription factor subunit NF-YB1, leafy cotyledon 1, endosperm-specific gene 41, Nuclear Factor YB1, NUCLEAR FACTOR-Y subunit B1, NUCLEAR FACTOR-Y subunit NF-YB1, NF-YB subunit 1, NF-YB family 1 | grain length (TO:0000734)<br>seed size (TO:0000391)<br>amylose content (TO:0000196)<br>seed anatomy and morphology trait (TO:0000184)<br>seed maturation (TO:0002661)<br>gel consistency (TO:0000211)<br>grain size (TO:0000397)<br>salt tolerance (TO:0006001)<br>auxin content (TO:0002672)<br>starch content (TO:0000696)<br>hot paste viscosity (TO:0000408)<br>vivipary (TO:0000619)<br>peak viscosity (TO:0000409)<br>seed dormancy (TO:0000253)<br>abscisic acid sensitivity (TO:0000615)<br>chalky endosperm (TO:0000266)<br>fat and essential oil content (TO:0000604)<br>1000-seed weight (TO:0000382)<br>seed development trait (TO:0000653)<br>cool paste viscosity (TO:0000379)<br>seed quality (TO:0000162)<br>gelatinization temperature (TO:0000462)<br>grain thickness (TO:0000399) | seed maturation stage (PO:0007632)<br>central endosperm (PO:0006220)<br>endosperm (PO:0009089)<br>aleurone layer (PO:0005360)<br>seed development stage (PO:0001170)<br>endosperm development stage (PO:0007633) | Masiero S et al. J Biol Chem, 277(29):26429-35 (2002) [PubMed]<br>Thirumurugan T et al. Mol Genet Genomics, 279(3):279-89 (2008) [PubMed]<br>Sun X et al. Gene, 551(2):214-21 (2014) [PubMed] | 食味・品質 |
| Os06g0133000 | Os06t0133000-01 | chr06:1765622..1770574 | WX1, Wx, wx (Wx(am)), WX-B, GBSS-I, GBSS, OsGBSSI, GBSSI, GBSS1, OsGBSS1, GSS, OsWx | GLUTINOUS ENDOSPERM, waxy, Waxy, glutinous endosperm, WAXY, "Granule-bound starch synthase, "Granule-bound starch synthase 1, chloroplastic/amyloplastic", Granule-bound starch synthase I, UDP-glycogen synthase, chloroplast precursor", glycogen [starch] synthase, Granule-bound glycogen synthase, UDPG-glycogen transglucosylase, uridine diphosphoglucose-glycogen glucosyltransferase, glycogen [starch] synthetase, Granule-bound glycogen [starch] synthase, UDPG-glycogen synthetase, granule bound starch synthase I, granule-bound starch synthase 1, granule-bound starch synthase I | amylose content (TO:0000196)<br>inflorescence development trait (TO:0000621)<br>fruit flavor trait (TO:0002694)<br>chalky endosperm (TO:0000266)<br>glutinous endosperm (TO:0000098)<br>submergence sensitivity (TO:0000286)<br>1000-seed weight (TO:0000382)<br>salt tolerance (TO:0006001)<br>nitrogen sensitivity (TO:0000011)<br>seed development trait (TO:0000653)<br>seed quality (TO:0000162)<br>water stress trait (TO:0000237)<br>heat tolerance (TO:0000259)<br>starch content (TO:0000696) | endosperm (PO:0009089)<br>seed imbibition stage (PO:0007022)<br>inflorescence development stage (PO:0001083)<br>seed (PO:0009010)<br>0 seed germination stage (PO:0007057)<br>seed maturation stage (PO:0007632)<br>seed development stage (PO:0001170) | Wang ZY et al. Plant J, 7(4):613-22 (1995) [PubMed]<br>Zhao Q et al. Nat Genet, 50(2):278-284 (2018) [PubMed]<br>Yong-zhong Li et al. Plant Science, 108(2):181-190 (1995) [DOI] | 食味・品質 |

**Fig. S11** Agronomically important genes filtered by the keyword "amylose content”.
